# Missing Data and Technical Variability in Single-Cell RNA-Sequencing Experiments

**DOI:** 10.1101/025528

**Authors:** Stephanie C. Hicks, F. William Townes, Mingxiang Teng, Rafael A. Irizarry

## Abstract

Until recently, high-throughput gene expression technology, such as RNA-Sequencing (RNA-seq) required hundreds of thousands of cells to produce reliable measurements. Recent technical advances permit genome-wide gene expression measurement at the single-cell level. Single-cell RNA-Seq (scRNA-seq) is the most widely used and numerous publications are based on data produced with this technology. However, RNA-Seq and scRNA-seq data are markedly different. In particular, unlike RNA-Seq, the majority of reported expression levels in scRNA-seq are zeros, which could be either biologically-driven, genes not expressing RNA at the time of measurement, or technically-driven, gene expressing RNA, but not at a sufficient level to detected by sequencing technology. Another difference is that the proportion of genes reporting the expression level to be zero varies substantially across single cells compared to RNA-seq samples. However, it remains unclear to what extent this cell-to-cell variation is being driven by technical rather than biological variation. Furthermore, while systematic errors, including batch effects, have been widely reported as a major challenge in high-throughput technologies, these issues have received minimal attention in published studies based on scRNA-seq technology. Here, we use an assessment experiment to examine data from published studies and demonstrate that systematic errors can explain a substantial percentage of observed cell-to-cell expression variability. Specifically, we present evidence that some of these reported zeros are driven by technical variation by demonstrating that scRNA-seq produces more zeros than expected and that this bias is greater for lower expressed genes. In addition, this missing data problem is exacerbated by the fact that this technical variation varies cell-to-cell. Then, we show how this technical cell-to-cell variability can be confused with novel biological results. Finally, we demonstrate and discuss how batch-effects and confounded experiments can intensify the problem.

## 1. Introduction

Single-cell RNA-Sequencing (scRNA-seq) has become the primary tool for profiling the transcriptomes of hundreds or even thousands of individual cells in parallel. In contrast to the standard RNA-seq approach, which is applied to samples containing hundreds of thousands of cells and therefore measures average gene expression level across cells, scRNA-seq measures gene expression in a single cell. To distinguish these two technologies we refer to the latter as *bulk RNA-seq*. Today scRNA-seq is increasingly being used across a diverse set of biomedical applications such as profiling the transcriptomes of differentiated cell types^1-4^, profiling the changes in cell states^5, 6^, identifying allele-specific expression^7, 8^, spatial reconstruction^9, 10^ and the classification of subtypes^11-13^.

While scRNA-seq data provides a new level of data resolution, it also results in a larger number of genes reporting the expression level to be zero, or practically zero, as compared to using bulk RNA-seq^14^. A gene reporting the expression level to be zero can arise in two ways: (1) the gene was not expressing any RNA at the time the cell was experimentally isolated and processed prior sequencing (referred to as structural zeros^15^) or (2) the gene was expressing RNA in the cell at the time of isolation, but not at a sufficient level to be detected in the experimental procedure to capture and process the RNA prior to sequencing (referred to as dropouts^15, 16^). While the former is a type of biological event, the latter is purely technical as it stems from the limitations of current experimental protocols to detect low amounts of RNA in a cell, referred to as capture efficiency^17^.

Batch effects are commonly found in high-throughput data^18^ and given the way that scRNA-seq experiments are conducted, there is much room for concern regarding confounding^18^. Specifically, batch effects in scRNA-seq experiments occur when cells from one biological group or condition are cultured, captured and sequenced separate from cells in a second condition (Figure S1). However, due to the nature of certain experimental scRNA-seq protocols, which restrict the way cells are captured and sequenced separately, sometimes standard balanced experimental designs are not possible^14, 18-20^. This reality makes it particularly important to be cautious about the potential for correlated variability induced by technical factors.

The unwanted variability introduced by batch effects can be particularly troublesome in scRNA-seq data because one of the most common applications has been the use unsupervised learning methods^21-26^ such as data exploration after dimensionality reduction or clustering to identify novel or rare subpopulations of cells^11-13^. Although a diverse set of techniques are used in these papers, both linear dimensionality reduction techniques, such as principal component analysis (PCA)^21^, and non-linear ones, such as t-Stochastic Neighbor Embedding (t-SNE)^26^, rely on computing distances between the cell expression profiles. Given that the majority of genes in a cell report the expression level to be zero and that the proportion of zeros varies greatly from cell to cell, it is not surprising that the distance estimates between cells are greatly influenced by the proportion of zeros^27, 28^. However, it remains unclear to what extent this cell-to-cell variation is being driven by technical rather than biological variation.

We begin this article by describing the publicly available scRNA-seq data sets we used, which includes studies with only scRNA-seq data and studies with scRNA-seq and a matched bulk RNA-seq sample measured on the same population of cells. In the next section, we survey a large number of published scRNA-seq studies and illustrate the wide range of variation in the proportion of genes reporting the expression level to be zero across cells and studies (Section 3.1). Then, we present evidence that some of these reported zeros are driven by technical variation by demonstrating that scRNA-seq produces more zeros than expected and that this bias is greater for lower expressed genes (Section 3.2). In addition, we show that the consequences of this missing data problem are exacerbated by the fact that the technical variation of the probability of a gene being detected varies from cell to cell. Then, we illustrate that the proportion of genes reporting the expression level to be zero is a major source of cell-to-cell variation and this variability is partly driven by a mathematical artifact related to the transforming data in the original scale, but computing distances in the log scale (Section 3.3). Finally, we consider several case studies showing how differences in the detection rates can be driven by batch effects, which in turn can result in the false discovery of new groups (Section 3.4).

## 2. Data Description

A scRNA-seq experiment typically involves randomly sampling and capturing single cells from a population of cells, isolating the mRNA from the individual cells, reverse transcribing the RNA into cDNA, and sequencing the cDNA using massively parallel sequencing technologies^19, 29, 30^. Strengths and weaknesses of different scRNA-seq experimental protocols vary^31-33^ in the cost per cell, the sensitivity to capture and convert RNA to cDNA and the accuracy to quantify the concentration of RNA, leading to differences in the number of cells sequenced per study and the number of features detected per cell. This experimental process is particularly challenging and laboratory protocols are still under intense development.

### 2.1 scRNA-seq data sets

We examine fifteen publicly available scRNA-seq data sets that included at least 200 samples with preprocessed and normalized expression data available on GEO^34^ (Table 1). These data sets were created using six different scRNA-seq protocols for sequencing^1, 12, 35-39^ and five studies include the use of unique molecular identifiers^40^ (UMIs) for counting specific cDNA molecules. For the ten studies not using UMIs we examine the data as submitted to GEO, with one exception^11^. These ten studies reported measurements in either Transcripts per Million^41^ (TPM), Reads Per Kilobase of transcript per Million mapped reads^42^ (RPKM) or Fragments Per Kilobase of transcript per Million mapped reads^43^ (FPKM), so each sample was corrected for gene length and library size. The one exception^11^ uploaded data that was de-trended so that measurements for each gene averaged to zero across cells. For this particular study, we downloaded the raw sequencing files data from the Sequence Read Archive (SRA)^44^ and computed expression in TPM units using Kallisto^45^. In the studies that used UMIs for molecule counting^1, 3, 9, 12, 46^, the data uploaded to GEO was not normalized for library size, so to assure that these data were in similar units to the rest of our studies, we followed a published procedure^1^ that normalizes each gene or transcript count by dividing by the total number of UMIs per cell and multiplies by a scaling factor (10^6^). We refer to this unit as Counts Per Million (CPM).

**Table 1:** Description of processed single-cell RNA-Seq data sets. Column 1 shows the publications. Column 2 shows the organism. Column 3 shows the single-cell technology used for sequencing. Column 4 shows the number of cells (samples) included in the study. Column 5 shows the number of genes included in the data uploaded to the public repository. Column 6 indicates the units in which the values were reported. Column 7 shows the level of confounding between biological condition and batch effect quantified using the standardized Pearson contingency coefficient as a measure of association. The percentage ranges from 0% (no confounding) to 100% (completely confounded). Column 8 provides the citation for the study. + The main purpose of this study was to investigate monoallelic gene expression in mouse embryos, but here we consider the different developmental stages (oocyte to blastocyst) as the biological condition as an example.

| Study | Organism | Single-cell RNA-Seq protocol | Number of cells | Number of genes or transcripts | Processed data available | Confounding (%): | Ref |
| --- | --- | --- | --- | --- | --- | --- | --- |
| Deng et al. (2014) | Mouse | SMART-Seq | 286 | 22,958 | RPKM | 96.6<sup>+</sup> | <sup>8</sup> |
| Guo et al. (2015) | Human | Tang et al. (2009) | 154 | 23,394 | FPKM | 82.1 | <sup>55</sup> |
| Kowalczyk et al. (2015) | Mouse | SMART-Seq | 533 | 8,422 | TPM | 84.8 | <sup>58</sup> |
| Kumar et al. (2014) | Mouse | SMART-Seq | 361 | 22,443 | TPM | 97.1 | <sup>56</sup> |
| Patel et al. (2014) | Human | SMART-Seq | 430 | 5,948 | TPM | 98.9 | <sup>11</sup> |
| Treutlein et al. (2014) | Mouse | SMART-Seq | 198 | 23,745 | FPKM | 92.8 | <sup>2</sup> |
| Shalek et al. (2014) | Mouse | SMART-Seq | 383 | 27,723 | TPM | 100 | <sup>6</sup> |
| Trapnell et al. (2014) | Human | SMART-Seq | 306 | 47,192 | FPKM | 100 | <sup>5</sup> |
| Burns et al. (2015) | Mouse | SMART-Seq | 249 | 26,585 | TPM | NA | <sup>57</sup> |
| Leng et al. (2015) | Human | SMART-Seq | 458 | 19,804 | TPM | NA | <sup>65</sup> |
| Bose et al. (2015) | Human | CEL-Seq | 247 | 17,450 | UMI | NA | <sup>46</sup> |
| Jaitin et al. (2014) | Mouse | MARS-Seq | 4,466 | 20,190 | UMI | NA | <sup>12</sup> |
| Macosko et al. (2015) | Mouse | Drop-Seq | 49,300 | 16,961 | UMI | NA | <sup>1</sup> |
| Satija et al. (2015) | Zebrafish | SMART-Seq | 1,152 | 13,902 | UMI | NA | <sup>9</sup> |
| Zeisel et al. (2015) | Mouse | STRT-Seq | 3,004 | 19,972 | UMI | NA | <sup>3</sup> |

Although details of the experimental protocols, which can help define groupings that may lead to technical batch effects, are not always included in the annotations that are publicly available, one can extract informative variables from the raw sequencing (FASTQ) files^47^. Namely, the sequencing instrument used, the run number from the instrument and the flow cell lane. Although the sequencing is unlikely to be a major source of unwanted variability, it serves as a surrogate for other experimental procedures that very likely do have an effect, such as the starting amount of RNA in a cell, capture efficiency, PCR amplification reagents/conditions, and cell cycle stage of the cells^6, 48-50^. Here we will refer to the resulting differences induced by different groupings of these sources of variability as *batches.*

### 2.2 scRNA-seq data sets with matched bulk RNA-seq data

To help determine if the increased proportion of zero in scRNA-seq is explained by biology or technical biases, we examined three publicly available scRNA-seq data sets^5, 51, 52^ that included a matched bulk RNA-seq sample measured on the same population of cells with preprocessed and normalized expression data available on GEO. One of these studies^5^ is one of the 15 studies described in the previous subsection. The other two studies^51, 52^ sequenced only 18 and 96 cells, respectively, thus were not included in the 15 large studies.

## 3. Results

### 3.1 The proportion of reported zeros varies from cell to cell and from study to study

We define the *detection rate* as the proportion of genes in a cell reporting the expression levels greater than a predetermined threshold δ. In this paper, we used δ =1 based on exploratory data analysis as this revealed two clear modes in the gene expression distribution (Figure S2), which we interpreted to be associated with background noise and signal respectively, with the lower mode defined as values below or equal to a TPM, FPKM, RPKM or CPM threshold of δ =1. This threshold has been previously used by Shalek et al. (2014)^6^ and accommodates the bimodality^6, 16, 51, 53, 54^ of scRNA-seq data that is not found in bulk RNA-Seq. We found wide variation in the detection rate across cells in all studies: from < 1% detected to 65% (Figure 1). Similar results were obtained if we set δ =0 (Figure S3). For studies including groups known to have different gene expression profiles, we stratified by biological group to minimize the possibility of a biological explanation and also found wide variation (Figures S4-S9). We also note the detection rate is not necessarily dependent on sequencing depth (Figure S10).

**Figure 1:**
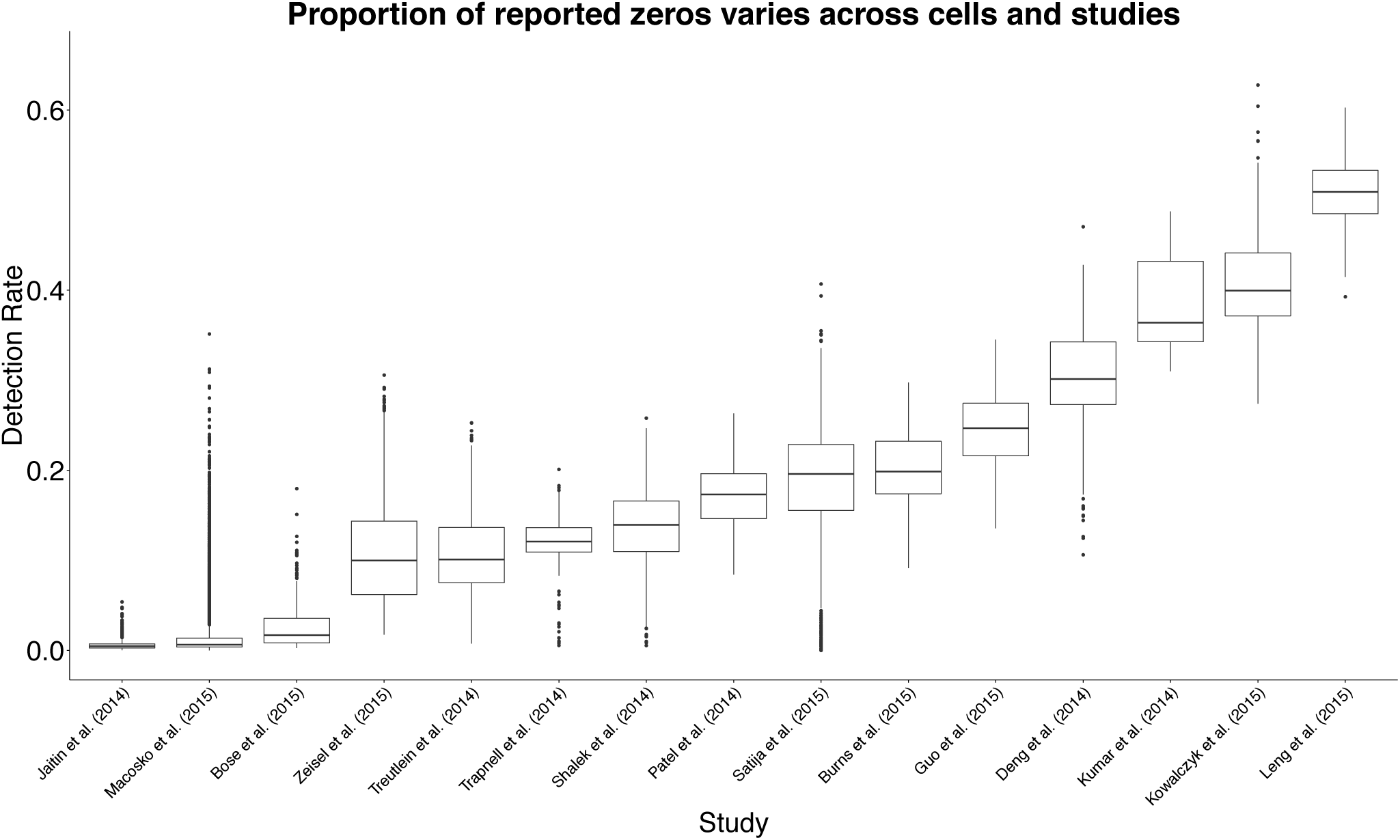
Boxplots of the *detection rate*, or the proportion of genes in a cell reporting expression values greater δ =1 calculated for each cell across fifteen publicly available scRNA-seq studies. The detection rate across cells and studies ranges from less than 1% to 65%.

### 3.2 scRNA-seq data contains more zeros than what is expected from biological variation

To demonstrate that there are more zeros in scRNA-seq data than what is expected from biological variation, we examined the gene expression of cells measured both on scRNA-seq and bulk RNA-seq. We found a bias consistent with a technical explanation. The details follow.

Denote the expression level for the *g*^*th*^ gene and *i*^*th*^ cell as *x*_*gi*_ where *i* = 1, …, *n*. The expression for the *g*^*th*^ gene in bulk tissue composed of these cells will then be:

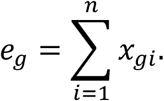

Bulk RNA-seq produces an estimate proportional to this quantity and includes measurement error (*ε*_*g*_):

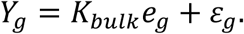

Here *K*_*bulk*_ is a normalizing constant needed to account for the fact that experimental protocols and normalization procedures are adjusted to assure that the average or sum of measurements from each experiment are approximately the same. Since a tissue sample will have millions of cells, we consider *n* to be large enough to be treated as infinity. Note that some of these *x*_*gi*_ can be zero even when *e*_*g*_ is a large number. In fact, this is part of the biological explanation for why single cell measurements have more zeros: an gene appearing expressed in bulk RNA-seq need not be expressed in every single cell at the time the cells were isolated and measured.

In a single cell experiment, we take a random sample of *N* cells from the population. We denote the expression values for these as *X*_*gi*_ where *i* = 1, …, *N*. Using scRNA-seq technology, we obtain measurements:

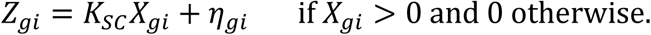

Here *K*_*sc*_ is the normalizing constant and *η*_*gi*_ is measurement error. Because the single cell data is a random sample, it follows that

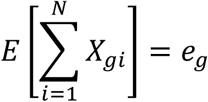

and therefore, if there is no biased induced from dropouts,

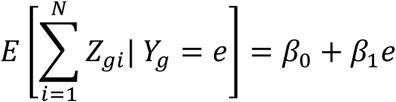

is a linear function with *β*_0_ and *β*_1_determined by the normalization constants and the variance of the measurement error, which has been reported to be relatively low. While in a typical scRNA-seq experiment, bulk RNA-seq measurements from the same tissue is not available, the three studies described in Section 2.2 with both bulk RNA-seq and scRNA-Seq from the same biological specimens permits us to check if this relationship holds.

As evidence that scRNA-Seq technology is working as expected, previous publications plot 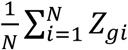 versus *Y*_*g*_ for each gene to show it generally follows a linear relationship with reported correlations around 0.80 (see, for example, Figure 1C in Shalek et al. (2013)^51^ which we reproduced in Figure 2A). However, a closer look at this plot reveals a problem: the linear relationship does not appear to hold for lowly expressed genes (Figure 2B). This same pattern is observed in the other two studies with bulk and scRNA-seq data (Figure S11).

**Figure 2:**
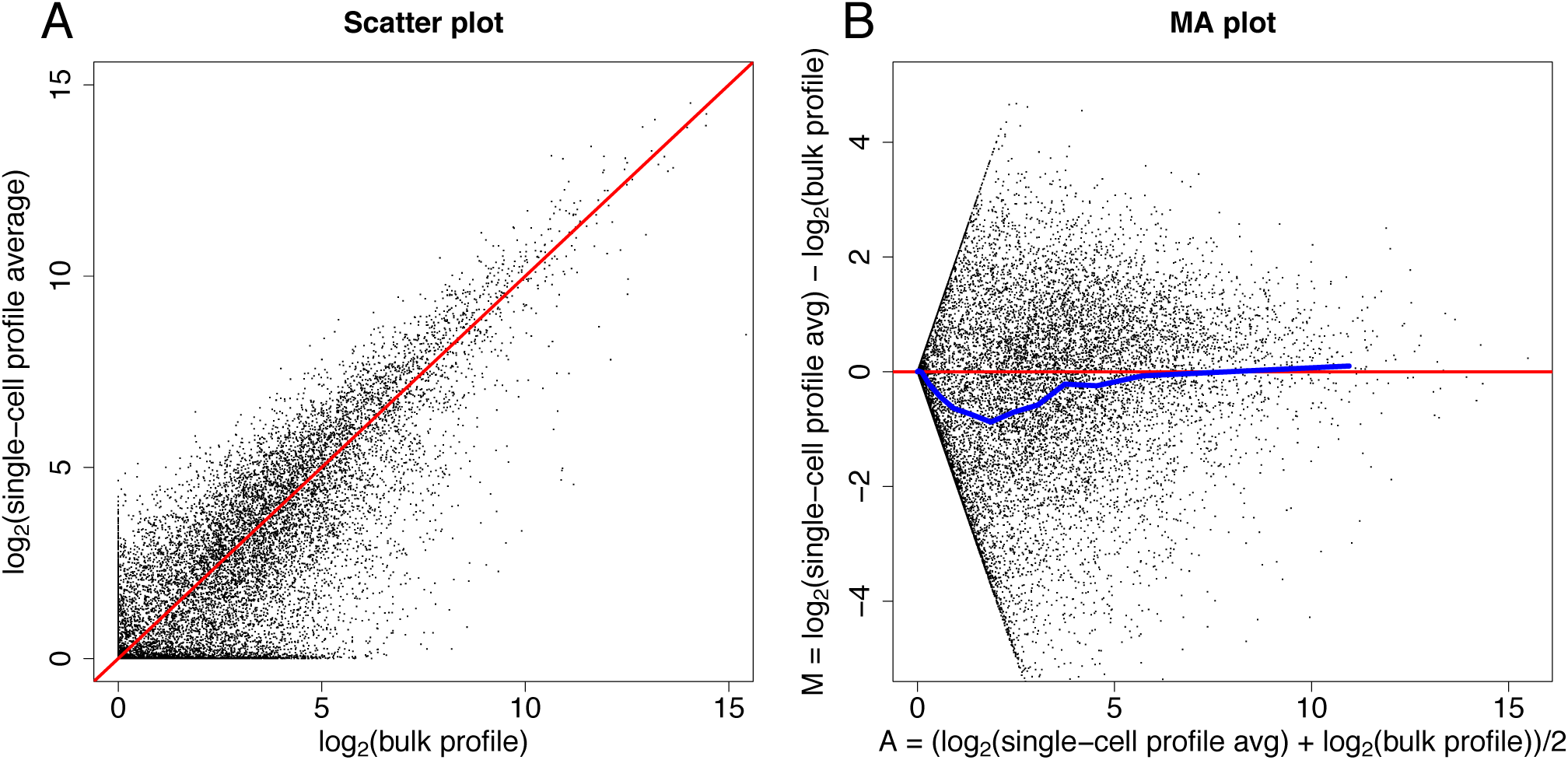
RNA-seq profiles compared to averaged scRNA-seq profile (**A**) Scatter plot comparing a bulk RNA-seq profile and an averaged scRNA-seq profile, which we reproduced from Figure 1C in Shalek et al. (2013)^51^. (**B**) The MA plot demonstrates there is a bias between the bulk profile and the single cell profile averaged across cells as the single cell profile averaged across cells is smaller than the bulk profile for low expressed genes.

To further explore this apparent bias, we stratified the values of *Y*_*g*_ and estimated the conditional expectations of 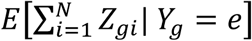 by averaging the scRNA-seq data in each strata. Plotting these against each other revealed a bias that increases as *e* becomes closer to zero (Figure 3). These results are very much consistent with the theory that some of the observed zeros are due to technical and not biological differences with the actual relationship being:

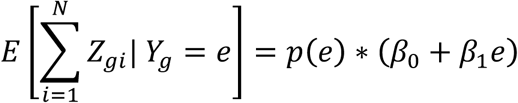

with *p*(*e*) the probability of a gene with expression *e* being detected. A crude estimate of *p*(*e*) can be obtained by

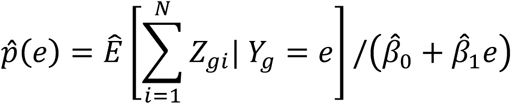

**Figure 3:**
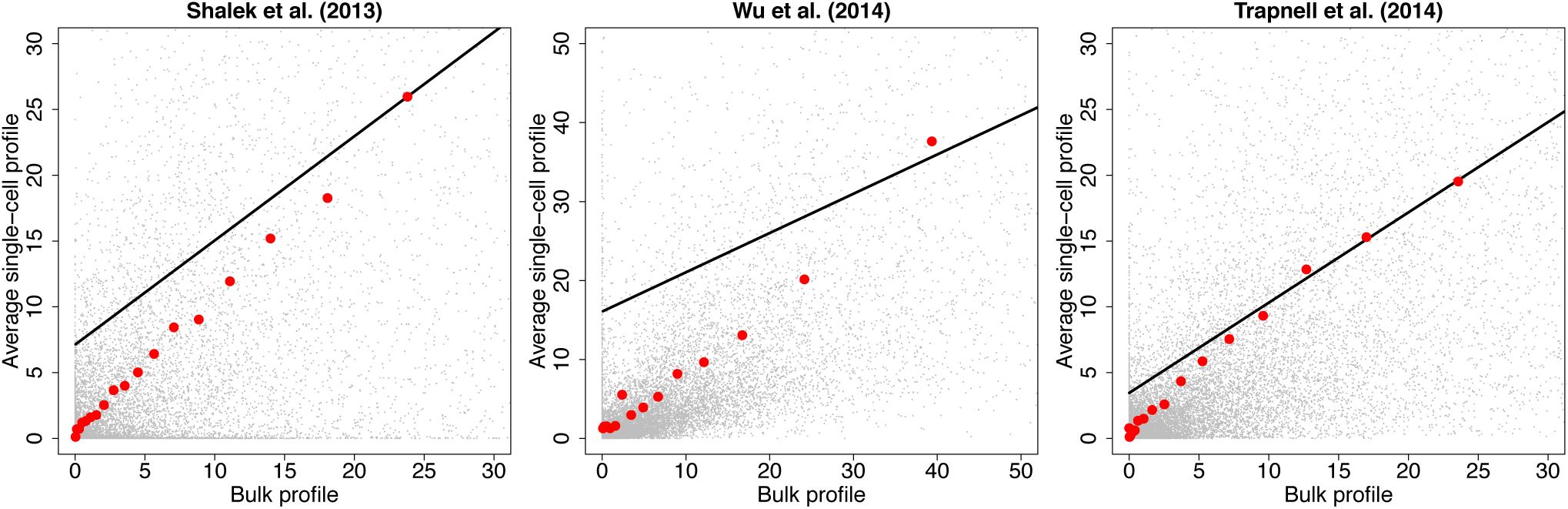
Plots comparing bulk and averaged scRNA-seq profiles that demonstrate evidence of more zeros in in scRNA-seq data for low expressed genes than what is expected. Data was obtained from three publicly available scRNA-seq studies that included a matched bulk RNA-seq sample measured on the same population of cells^5, 51, 52^. The red points are averages of the single cell profiles computed in strata defined by the bulk RNA-seq values. The black solid line is what we expect if there is no bias.

This estimate suggests *p*(*e*) follows a logistic function (Figure S12) as others have previously noted^16^. In other words, genes with a lower expression *e* are less likely to be detected, which suggests the zeros can be considered missing not at random as the probability of the missing value depends on the level of expression.

Motivated by Figure S12 we fit a logistic curve to determine the relationship between *Z*_*gi*_ > δ and *Y*_*g*_ for each cell *i*. We found that the biases induced by this missing data problem is exacerbated by the fact that the probability of a gene being detected varies cell to cell, as the estimates for the logistic curve’s intercept parameter are highly related to the detection rate (Figure S13). We also note that the slope estimates are between 0.53 and 1.31. For example, the slopes estimated using the Trapnell et al. (2014)^5^ data has an average of 0.82 and a standard deviation of 0.6 demonstrating the strong effect overall expression has on detection: note for example that a slope of 0.82 means that if the expression level is cut in half, the detection odds decrease by more than two fold since *e*^0.82^= 2.27.

### 3.3 Detection rate is a major source of cell-to-cell variation

Finak et al. (2015)^53^ showed that detection rates correlate with the first two principal components (PCs) in two scRNA-seq data sets^6, 53^. We confirmed this relationship on the fifteen publicly available scRNA-seq datasets we studied (Figures S14-S15). From this strong correlation it follows that estimated distances between cells are affected by differences in detection rate. We note that for five of these studies^3, 8, 55-57^, the primary variation along the first two principal components was correlated strongly with the biological groups known to have different gene expression patterns. For these studies, we stratified the data into these groups and found the same strong relationship between detection rates and first principal component. In this section, we present results that demonstrate that (1) this variability is partly driven by a mathematical artifact related to scaling the original data but computing distances in the log transformed data and that (2) that differences in detection rate can be completely driven by technical reasons which can in turn result in false discoveries.

Currently the most widely used unit for reporting expression values is the Transcripts per Million (TPM) unit. Using this unit guarantees that the sum of gene expression measurements are constant. This is also true for CPM and approximately true with the RPKM^42^ and FPKM^43^ units. However, distance calculations are performed after log transformations and cell expression profiles are not always reported as being centered (centered by removing overall mean expression from the *i*^*th*^ cell) in published analyses^1, 2, 57, 58^. We can show, mathematically, that if we normalize expression profiles to have the same mean across cells, the mean after the *f* (*x*) = log (*x* + *c*) transformation used for RNA-Seq data will not be the same and it will depend on the detection rate (Figure S16).

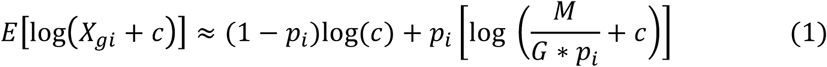

where *X*_*gi*_ is the expression value for the *g*^*th*^ gene and *i*^*th*^ cell, *c* is a pseudo count, *G* is the number of genes (or features), *M* is sequencing depth, and *p*_*i*_ is the marginal probability of detection for the *i*^*th*^ cell (mathematical details are provided in the Supplement). The implication of this result is that although the means are constant using across cells *i*, the means of the log-transformed data depend on the detection rate. In fact, when the sequencing depth is large, the mathematical relationship above is approximately a linear function of the detection rate. Because these mean values affect the entire vector, they can result in large overall variability and therefore be correlated with the first principal component PC. Therefore, the result in Figures S14-S15 can be explained by differences in mean values that correlated with the detection rate.

Not surprisingly, if we center the data before computing the PCs, then the correlation between the detection rate and the first PC decreases. However, despite the decrease, even after the centering the correlation is strong (Figures S17-S18). This also is not surprising given that not only the mean expression depends of the detection rate, but also the entire distribution of the non-zero measurements using (Figure 4).

**Figure 4:**
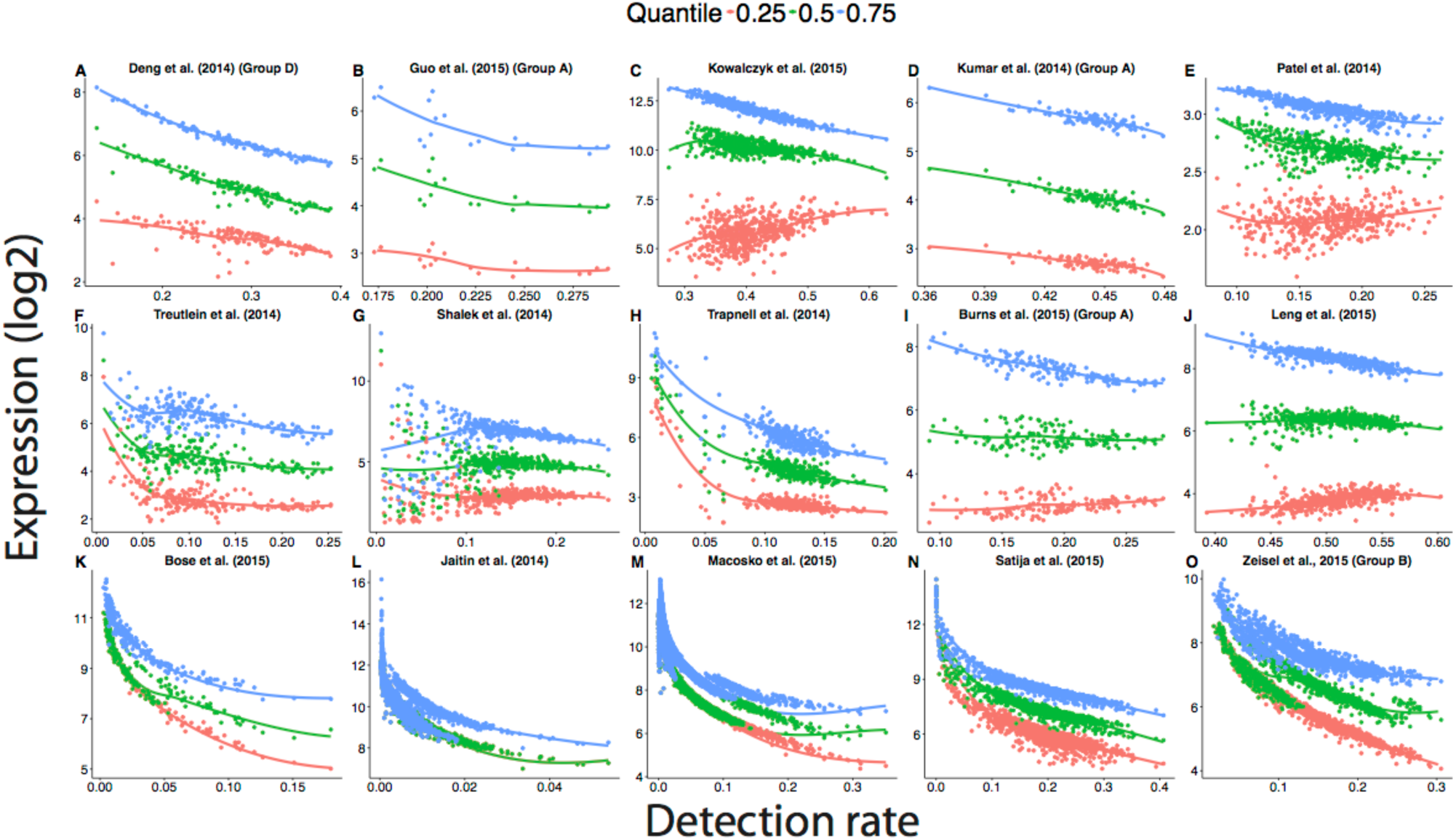
The distribution of gene expression changes with the detection rate using processed scRNA-seq data available on GEO. Failure to account for differences of the proportion of detected genes between cells over-inflates the gene expression estimates of cells with a low detection rate. The curves were obtained by fitting a locally weighted scatter plot smoothing (loess) with a degree of 1. Because the range of detection rate varied from study to study, the range of the x-axis differs across plots.

To demonstrate that technical variability can lead to differences in detection rates, which in turn can lead to false discoveries, we used a dataset composed of a group of cells from the same biological specimen, but processed at different times. Specifically, we used a subset of 118 single cells data from Patel et al. (2014), which were isolated from one tumor, but processed in two different sequencing instruments^11^. For these data, there are no biological reasons for the two groups, defined by the sequencing instrument used, to be different since the cells were randomly selected from the same tumor. If we apply an unsupervised clustering algorithm to these data, two clusters strong clusters appear (Figure S19) even after removing the cell mean before computing distances. A PCA plot shows that a batch effect drives the clustering (Figure 5A). We then note that the first PC strongly correlates with the detection rate (Figure 5B), which is substantially different between the two batches (Figure 5C). In addition, the differences in detection rates are highly related to the logistic curve’s intercept parameter when estimating the probability of a gene being detected, which varies cell to cell (Figure S20). Therefore, we see how a batch effect can produce differences in detection rate that drive distances between transcription profiles and leads to false discoveries.

**Figure 5:**
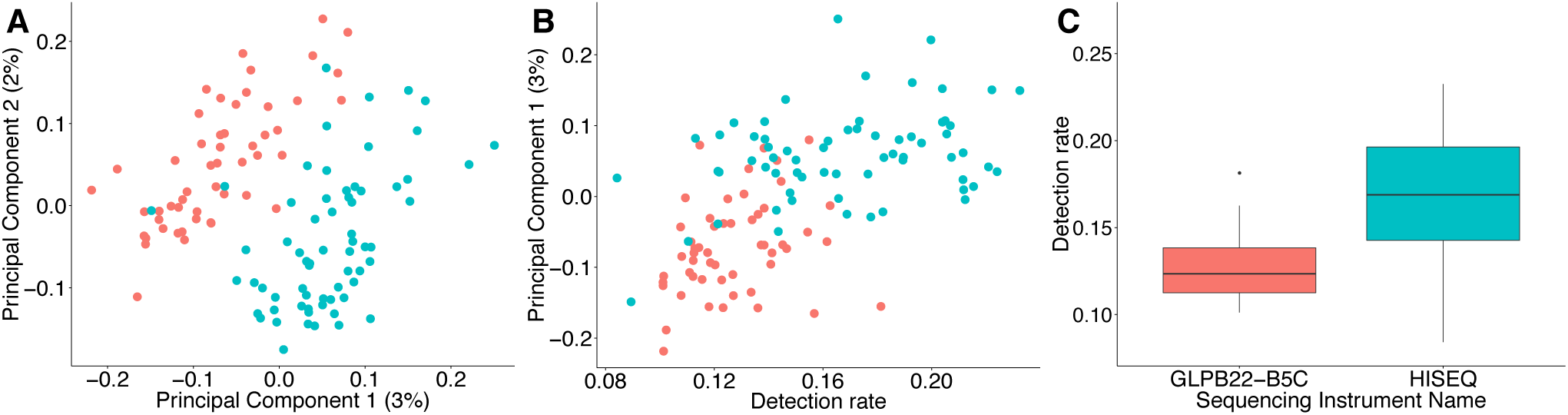
Illustration of how technical variation can lead to differences in detection rates, which in turn can lead to false differences. (A) Boxplots of detection rates from cells stratified by sequencing instrument used to sequence cells. (B) Using principal components analysis, scRNA-seq samples cluster by sequencing instrument. (C) Detection rate is associated with the first principal component.

### 3.4 The impact of the detection rate in applications of unsupervised learning methods

In the previous sections, we demonstrated how the detection rate is a major source of observed cell-to-cell variation, which can be driven by technical variation. For example, we considered cells from the same biological specimen for which we knew no differences should be discovered, but we found a batch effect that had differences in the detection rate and could drive the variation between the cells. In this section, we examine detection rates in data used in published studies as evidence for the discovery of new cell types.

In the first case-study^57^, we found differences in the detection rate between two groups of discovered cell types (Figure S21A), which was associated with how the cells were processed in different batches (Figure S21B) (Fisher’s Exact test: *p* = 0.002). For example, 12 out of 13 cells from one discovered cell types were processed in the same batch (Figure S21C), which had a smaller median detection rate than the other batches. Furthermore, the detection rate was associated with the first principal component (Figure S21D), which could be partly driving the variation across the two groups of discovered cell types. Similarly, we found differences in the detection rate between groups of discovered cell types in another other study^3^ (Figure S22), which was associated with how the cells were processed in different batches (Chi-squared test: *p* < 0.001).

## 4. Discussion

We have demonstrated how detection varies substantially across scRNA-seq experiments. We presented evidence that part of this variability is technically driven. Given the logistics of how scRNA-Seq experiments are performed and that fact that this technology is being used to discover new cell-types, batch effects are of particular concern. Specifically, when two groups of cells are cultured, captured and sequenced separately from another group of cells in a second condition, correlated measurements may lead to the incorrect conclusion that these groups have different expression profiles. This experimental limitation presents a challenge in distinguishing biologically driven differences from technical ones because it is logistically difficult to avoid processing cells from different biological specimens in different batches. For the eight studies that were interested in comparing predefined biological groups, we used the standardized Pearson contingency coefficient to assess the level of confounding between the run number from the sequencing instrument and outcome of interest and found values ranging from 82.1% to 100% (perfect confounding) (Table 1; Tables S1-S8). Note that with this level of confounding it difficult to impossible to parse technical from biological variation.

Furthermore, explicitly modeling confounding factors as in published batch correction methods^59-61^ is not appropriate in this context because the biological variation or signal of interest is often confounded with the unwanted technical variability. For the specific application of differential expression, a proposed solution is to account for differences in the proportion of detected genes by explicitly including it as a covariate in a linear regression model^53^. However, given the current levels of confounding, this approach will not be able to distinguish biological from technical effects. For example, some studies have demonstrated cells with different biological phenotypes can express a different number of genes^62^.

An experimental design solution is to use biological replicates, namely independently repeating the experiment multiple times for each biological condition (Figure S1). This approach allows for multiple batches of cells to be randomized across sequencing runs, flow cells and lanes as in bulk RNA-Seq. With this design we can then model and adjust for batch effects due to systematic experimental bias. A more detailed discussion of how these factors affect the experimental design has been recently published^14, 18, 19^.

## 5. Conclusions

Technical variability is considered to be a major challenge in the analysis of data measured on next-generation sequencing platforms. For example, amplification bias leading to batch effects has been shown to induce false positives in differential expression studies with bulk RNA-seq data^63, 64^. By examining three assessment experiments containing both bulk and single cell RNA-Seq data, we demonstrated that technical variability is a challenge in scRNA-Seq as well, with a major problem arising due to differences in capture inefficiencies. Using public data from fifteen studies, we showed that these inefficiencies lead to substantial differences in detection rates that lead to distortion in distance calculations, which in turn can lead to false discoveries when using unsupervised clustering.

## Acknowledgements

We thank Bradley Bernstein who provided comments that we used to improve the manuscript. This research was supported by NIH R01 grants GM083084, RR021967/GM103552 and HG005220 and NIH/NHGRI grant K99HG009007.

## Supplementary Material

Supplementary materials are available in a single PDF. All the code for this analysis is available on GitHub (https://github.com/stephaniehicks/scBatchPaper).

